# ImPartial: Partial Annotations for Cell Instance Segmentation

**DOI:** 10.1101/2021.01.20.427458

**Authors:** Natalia Martinez, Guillermo Sapiro, Allen Tannenbaum, Travis J. Hollmann, Saad Nadeem

## Abstract

Segmenting noisy multiplex spatial tissue images constitutes a challenging task, since the characteristics of both the noise and the biology being imaged differs significantly across tissues and modalities; this is compounded by the high monetary and time costs associated with manual annotations. It is therefore imperative to build algorithms that can accurately segment the noisy images based on a small number of annotations. Recently techniques to derive such an algorithm from a few scribbled annotations have been proposed, mostly relying on the refinement and estimation of pseudo-labels. Other techniques leverage the success of self-supervised denoising as a parallel task to potentially improve the segmentation objective when few annotations are available. In this paper, we propose a method that augments the segmentation objective via self-supervised multi-channel quantized imputation, meaning that each class of the segmentation objective can be characterized by a mixture of distributions. This approach leverages the observation that perfect pixel-wise reconstruction or denoising of the image is not needed for accurate segmentation, and introduces a self-supervised classification objective that better aligns with the overall segmentation goal. We demonstrate the superior performance of our approach for a variety of cancer datasets acquired with different highly-multiplexed imaging modalities in real clinical settings. Code for our method along with a benchmarking dataset is available at https://github.com/natalialmg/ImPartial.

## Introduction

Multiplex tissue imaging currently allows the user to detect several cellular and sub-cellular biologic signals within an intact tissue section. This technique allows for interrogation of spatial relationships, co-expression and derivative interactions. Currently, most techniques detect protein expression with antibody-based detection mechanisms, however, there are techniques that also allow RNA and DNA detection. Recent advances in these techniques have enabled the collection of large amounts of valuable data; this has outpaced manual labeling and highlights the need for automatic, semisupervised procedures to extract relevant information from these modalities. One task of interest is removal of background noise, which improves the joint visualization of the tissue across modalities. Another important task is the automatic segmentation of biological structures. Both of these tasks are challenging given the variability of noise, illumination, and biological structures that comes from different tissue types and acquisition modalities.

There are three predominant multiplex imaging signal detection modalities: (1) chromogenic, (2) fluorescence, and (3) mass spectrometry imaging. Chromogenic staining doesn’t allow visualization of isolated individual channels and hence is not used for staining more than 2-3 markers clinically (up to 8 markers can be stained) as individual signal delineation and overlap can be problematic with light microscopy. Fluorescence and mass spectrometry techniques allow for staining and visualization of individual channels and hence are the most prevalent multiplex imaging modalities, each having the potential to image over 40 biomarkers per tissue section. Denoising and instance segmentation of the individual channels/markers as well as composites within variant noise fields is a major barrier to automated algorithmic protein expression reporting in the clinical workspace. Current deep learning algorithms and commercial multiplex image analysis platforms often require preprocessing of images to partially denoise individual channels (via crude linearly filtering or k-nearest-neighbor (kNN) approaches (1)) as input for cell instance segmentation; when this partial denoising via linear filtering or kNN fails for the acquired images (e.g. Figure 1 top row), relevant data is removed from any further analysis (1,2) even though it could have been easily salvaged using for example our presented approach.

**Fig. 1.**
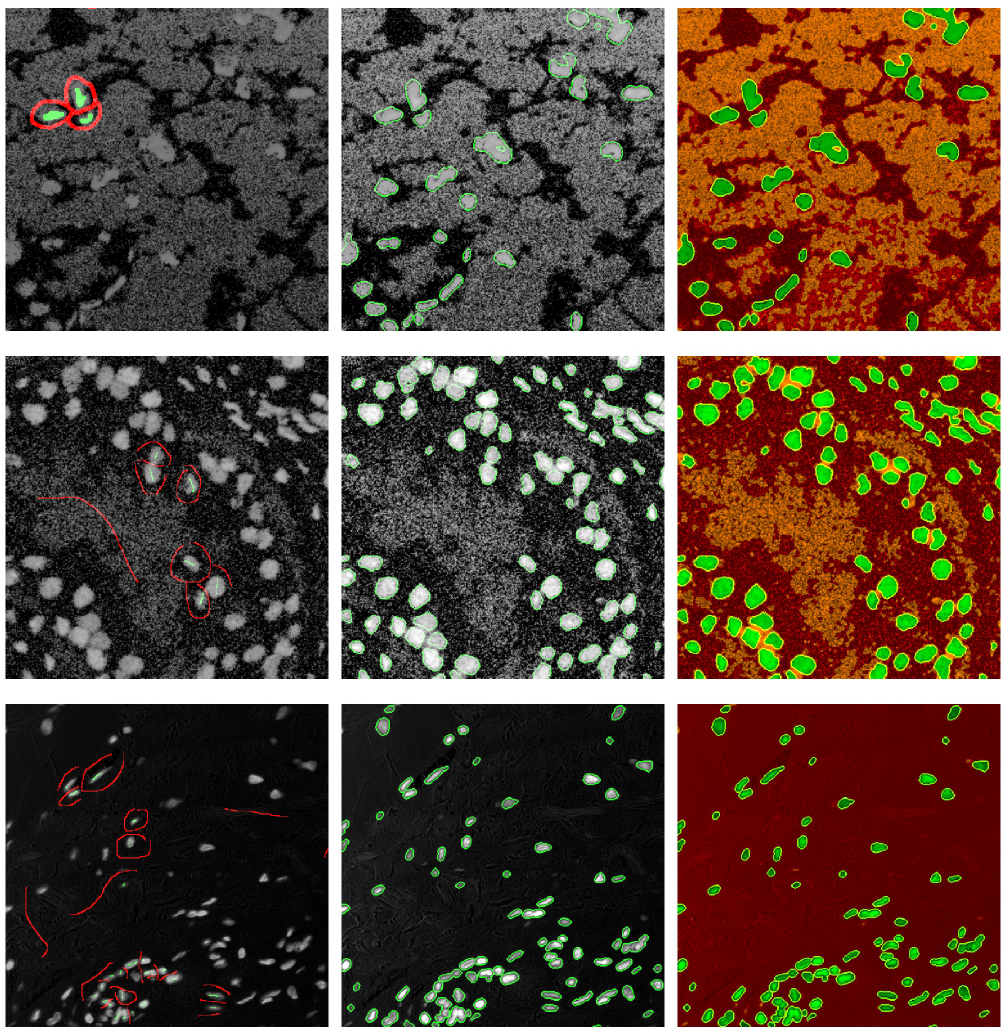
Here we show three examples, first and second row corresponds to MIBI image of breast tissue (dsDNA nuclei channel), third rows is a Vectra DAPI nuclei channel. The first column shows the input images and the annotated scribbles (background in red and foreground in green). The green contours in the images in the second column highlight the detected foregrounds, we modeled the cluster components as fixed-variance Gaussian distributions with parametric mean, a total of *M* = 3 clusters were used, with 2 of those modeling the background distribution, and 1 modeling foreground. Right images show the cluster assignments 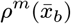 over the whole image, orange and red correspond to each background mode, and green indicates foreground.

Deep learning promises a solution for some of these problems, and has reached state-of-the-art-performance in segmentation of biomedical images (3–5), and, in particular, segmentation of nuclei and gland structures (6–10). However, these techniques require large amounts of ground-truth labels, a task that is labor intensive and needs significant time and monetary investment. Moreover, human annotations may suffer from inter-observer variability depending on the biological structure being segmented and the background noise. To cope with the aforementioned lack of annotations, different data augmentation methods have been proposed (11–15). The limitations of these methods lie in the assumption that the chosen augmentation family can adequately cover the full range of variation of the tissue to be segmented. Recent weakly supervised techniques, where only nuclei point annotations are provided, have been proposed to avoid reliance on data augmentation (16–18). These methodologies require fully annotated nuclei patches to generate coarse labels used to regularize a segmentation objective. Intermediate approaches such as (19) propose training a network that can assist in the segmentation process, simplifying the task of fully annotating new images. However, the latter method requires manual inputs (e.g. centroids) for all the cell instances in the image.

The use of scribbles is a viable alternative to full annotations due to its simplicity. A combination of scribbled-supervised learning and object dependent regularization has shown success in semantic segmentation of natural images (20–22). Scribble2Label (23) was recently proposed in the context of cell segmentation, outperforming methods such as (20, 24, 25). This method trains a network to provide a segmented image by iteratively estimating the unlabeled examples (pseudo-labels), but does not explicitly account for noise in the input images. Recently proposed deep learning self-supervised denoising techniques (26) have been shown to improve object segmentation when performed as a preprocessing step (27). Moreover, the recently proposed DenoiSeg (28) trains a U-Net to simultaneously output the segmented regions and a self-supervised denoised image (26). This approach does not require a large number of labeled images, and has shown promising performance on noisy microscopy data.

In this work, we present a method to identify instances of biological structures, e.g., cell nuclei. We propose a weakly-supervised scheme that uses a small number of scribbles, which are drawings indicating objects of interest and surrounding background or boundaries (see Figure 1), for training. Similar to DenoiSeg(28), we leverage the *blind-spot network* training scheme (26) to define a self-supervised auxiliary task. In our case, we define an image quantization objective that models the background and foreground as coming from a limited number of uni-modal distributions. This is a reasonable prior since in many microscopy imaging scenarios different objects appear at distinct intensity levels.

We evaluate our method on nuclei segmentation on a variety of single channel noisy mass spectrometry multiplexed ion beam images (MIBI) for breast, bladder and lung tissues. We also test our performance on cleaner acquisitions from Vectra fluoresence imaging modality. Finally, we test our method on two-channel input images (cytoplasmic and nuclear markers) to perform joint per-channel segmentation using a single neural network. We show how varying the number of scribbles affects the performance on partially annotated images as well as unseen ones. We compare against the recently proposed methods (23, 28) and a baseline model that implements the *blind-spot network* training scheme and only takes into account the segmentation loss on the available scribbles.

### Related Work

Using partial annotations instead of fully annotated images has been explored in the context of natural images (13, 20–22, 24, 25). Many of the weakly supervised techniques for nuclei and gland segmentation (16–19) require patches where partial labels are available for all the instances in order to boost the segmentation loss with coarse labels. Since in our approach we rely on having scribbles of a subset of the available instances in the image (e.g., nuclei), our work most closely relates to the recently proposed Scribble2Label (23). Scribble2Label assumes that partial scribbles for both foreground (instances) and background are provided. It generates and leverages pseudo-labels by iteratively refining the segmentation with high confidence estimations. In essence, this method first trains a network with only the available scribbles and then builds pseudo-label for the missing annotations by iteratively considering an exponential moving average over the pixels whose estimations were above a certain confidence threshold. Scribble2Label has shown better performance on cell segmentation than other weakly-supervised segmentation methods such as (20, 24, 25). They also show that there is no need for fully annotated data or a large amount of scribbles to have a good performance on different segmentation tasks, something that we also validate in this work. Scribble2Label does not particularly account for noisy observations other than on their confidence parameter threshold.

When dealing with segmentation for noisy images, applying a denoising step has been shown to be potentially beneficial (27). In this context, the DenoiSeg (28) framework proposes a way to simultaneously train a U-net that outputs self-supervised denoised images using the Noise2Void (26) objective plus background, foreground and border segmentation. This approach showed that having a network with a parallel denoising objective can improve the segmentation performance and reduce the number of annotated images. Moreover, it is shown that this was better than a two-stage approach alternative (27) (self supervised denoising and then segmentation). The technique trains against fully annotated image patches but can be made to work with scribbles as well. This method is closely related to ours since we also leverage the *blind spot network* from (26) to deal with noisy images. We instead choose to have a quantization (mixed regression and classification) loss as our side objective. Our approach is related to (29) where the Mumford-Shah functional is adapted to be used as a regularization in semi-supervised semantic segmentation with deep neural networks. Their approach, however, does not work well for noisy images.

## Method

In this section, we describe our method to segment multichannel noisy images when only a few scribbles are provided. Since the size and scale of the input images may vary across acquisitions, we use a U-net architecture to train a space-preserving image segmentation and classification network. For this, we assume that each segmentation class can be reasonably modeled as a sparse mixture of distributions. This sparsity assumption introduces quantization into our denoised image; the resulting reconstruction loss is used to improve the segmentation objective.

Assume we have access to a dataset containing *n* pairs of noisy images and scribbles 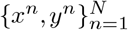. Here *x* ∊ **R**^*W×H×C*^ is a noisy input image and *y* ∊ {0,1}^*W×H×K*^ is the corresponding ground truth scribble per-pixel, indicating which of the *K* classes is present at the location. For notational compactness, we use the sub-index *i* to refer to the spatial location of a pixel instead of 2-coordinate location (*i,j*); we also overload the scribble notation *y_i_* where *y_i_* sometimes satisfies ‖*y_i_*‖_1_ = 1 (one-hot class encoding where scribbles are present), but otherwise satisfies ‖*y_i_*‖_1_ =0 to indicate no scribble is present. We first define the training losses for a single channel (*C* = 1) in what follows; the extension to multiple channels is described subsequently.

### Mixture Loss

Following the *blind-spot network* proposed in Noise2Void (26), for each image patch *x^b^* in our training batch *b ∊ B*, we generate a partial copy 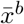, where a random set of pixels *I_b_* is substituted by random values in the vicinity 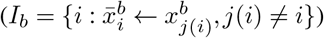. We then define the reconstruction loss of the batch for mixture model with *m ∊ M* components as follows:

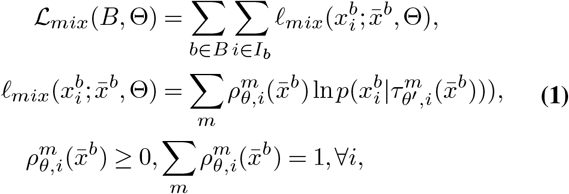

where 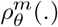 is a parametric (in *θ*) function that outputs the cluster membership per pixel, 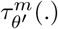 computes the sufficient static of the distribution of the *m*-th component. The dimensions of 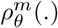 and 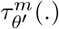 are therefore *W × H × M*, and *W × H × (M × S)* respectively, where *S* is the number of sufficient statistics for the cluster distribution (e.g., *S* =1 for a Gaussian distribution with fixed variance). Note that each pixel is modeled as coming from one of the cluster distributions, with cluster assignment probability 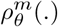, as opposed to modelling the pixel as a mixture of distributions, like would be the case for a Gaussian mixture model.

The components statistics 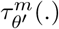 are a function of the observed patch 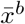, and therefore present a dynamic behavior that may be learned from data using a neural network. This allows the adjustment to non-homogeneous images (e.g., nonuniform intensity as in the example shown in Figure 1). In implementation details, we describe the additional assumptions we make to preserve consistency of this component mixture across patches. Both 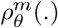 and 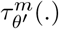 are implemented as output channels of our U-Net network, we use the notation © to refer to the entire network parameters, which are partly shared across *θ* and *θ’*.

### Scribble Loss

Given an image patch *x^b^* and its corresponding one-hot encoded scribbled image *y^b^*, we denote the set of annotated pixels of class *k ∊ K* as 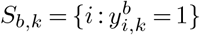. We assume that each class *k ∊ K* is associated with *m_k_* unique components of the mixture presented in Equation 1, and propose the following segmentation loss for each *k* ∊ *K*.

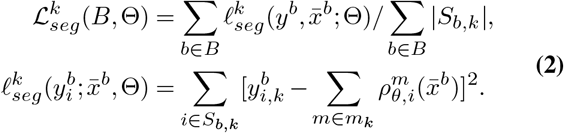

Note that this loss is equivalent to a class-balanced Brierscore loss on the pixels where class labels are available 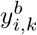, and their associated membership class 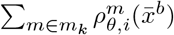 (computed as a sum over cluster labels); class balancing is achieved by dividing the per-class score over the number of annotated scribbles 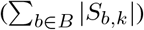. The combination of this objective with the mixture loss enforces regularity on the cluster assignments and therefore on the component statistics 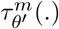. The dual perspective is that we encourage our recovered class labels to be consistent with the reconstruction objective defined for the mixture loss.

### Joint loss

We combine the quantization mixture loss and the scribble segmentation loss that makes use of the available scribbles into a single training loss. Given a batch of images of size *B*, we define the following joint objective:

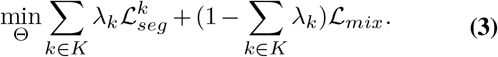

Note that here *λ_k_* represents the weight given to each segmentation class loss. We choose *λ_k_ >* 0 such that 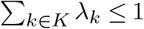. Since the reconstruction loss 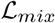 is a regularization task, we choose a small budget 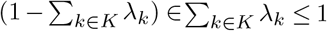. For all experiments, we set the value of *λ_k_* to be the same across all classes *k ∊ K*s.

### Implementation details

Here we describe the implementation of the proposed scheme, shown in Figure 2. Given an image patch, we apply the Noise2Void masking method to generate the blind spot patch that is observed by our network architecture, implemented as a U-Net. The output channels of the network comprised of *M* channels under a softmax nonlinearity, which generate the mixture mask *ρ^m^*(·), and an additional *M* channels outputting the sufficient statistics of the mixture components *τ^m^*(·) We chose fixed-variance Gaussian distributions to model each component (the joint estimation of mean and variance of the components did not meaningfully improve the segmentation objective). With these outputs, *ρ^m^*(·) and *τ^m^*(·), we compute the joint loss and propagate the gradients back to the network parameters.

**Fig. 2.**
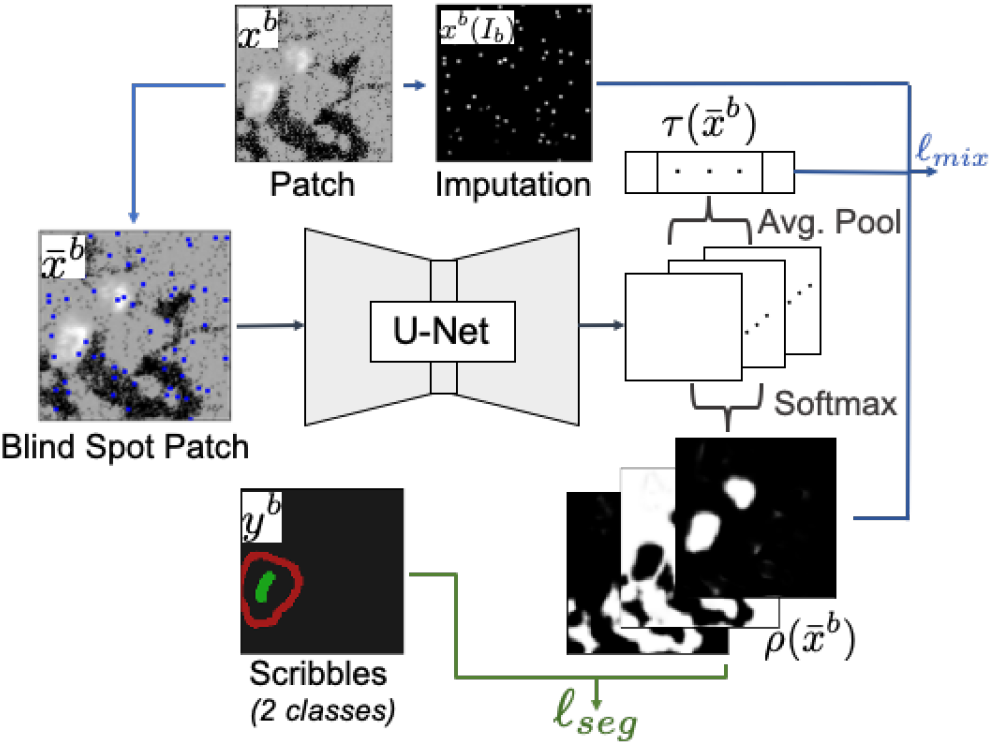
Overview of the processing pipeline. Each image patch is separated into an imputation patch and a blind spot patch. The blind spot patch is fed through the U-Net to recover the component mixture 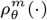 and the component statistics 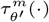, the latter statistics are averaged across the entire patch to enforce component consistency, both the component statistics and component mixture are used to compute the mixture loss for the patch. Simultaneously, a scribble containing a small number of ground truth segmentations for the patch is used to compute the scribble loss. Both losses propagate gradients back to the U-Net architecture on the backward pass.

We build our model on top of the frameworks proposed in (26, 28, 30), unless otherwise specified. We use a U-Net with depth 2, 64 initial feature maps, and convolution kernel size of 3. We use a batch size of size 64, trained for 100 epochs, where each epoch consisted of 50 batch descent iterations. We use Adam as our optimizer method with learning rate 4 x 10^-4^, stopping criteria was based on validation performance on joint loss. In all examples, the network converges before the 50-th epoch.

All examples shown in the paper contain two scribble classes (foreground/instances and background/boundaries). In all cases, we modeled foreground and background with two components each. The extra foreground component proved to be unnecessary in many cases, but did not affect performance when compared to a single-component foreground. Since our prior assumption is that foreground structures tend to present overall higher intensity levels than background, we explicitly codified this prior by adding the background component mean (mean of the background sufficient statistics) to the foreground sufficient statistic.

### Extension to multiple channels

We extend the framework to multiple channels (input image 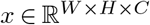) in the natural way by computing the mixture (quantization) statistics jointly across all channels *C*. Across all experiments, each component *M* in the mixture is modeled as a fixed-variance Gaussian with mean 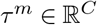. The component mask 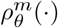 is therefore shared across channels. Since we assume the scribbled mask is related to objects that appear as foreground components on at least one channel, we can still compute the scribble mask by summing over the foreground components of the multi-channel mixture mask. The joint loss therefore remains unchanged.

In this section we provide a description of the scribble annotating scheme, and of the datasets used for evaluation. We worked with datasets acquired from the fluorescence imaging platform Vectra Polaris, and the mass spectrometry imaging platform MIBI. We then present the performance of competing methods for each described dataset.

### Annotating scribbles

In this work, we segmented cell nuclei and cytoplasm instances. We assume that the expert an-notator would not provide full pixel-wise segmentation of the objects of interest, but would instead draw scribbles to indicate the presence of foreground (instances) or background. For standard segmentation, we assume the foreground scribble contains an object of interest (e.g., a line across a nucleus of interest), and that additional scribbles for surrounding background are provided. For cell and cytoplasm segmentation, we assumed the pathologist would also draw a contour inside the cytoplasm.

Since we have access to fully annotated masks, we simulated the pathologist interaction as follows. We randomly select a field of view in the image (in our case a box of size 32 x 32, with no particular selection criteria) and provide scribbles for the instances and background available in the region. This simulates how an actual human annotator would interact with the software, since it is a less demanding workflow than randomly selecting an instance on the entire image for each new scribble. Foreground scribbles of nuclei instances were generated from their skeletonized masks. Background scribbles are the skeletonized mask of the background in the bounding box. For cytoplasm segmentation, an additional contour of the eroded instance mask was provided (this is because in most of the 2-channel composite examples, the skeletonized scribble of the nuclei and cytoplasm significantly overlapped with the one corresponding to just the nuclei). In our experiments, we vary the number of scribbles provided by repeating the above process until the scribble budget is depleted.

### Datasets

We tested our method on datasets acquired from fluorescence imaging platform, Vectra Polaris^1^ and mass spectrometry imaging platform, MIBI (Mulitplexed Ion Beam Imaging)^2^. Background signal in fluorescent imaging (Vectra) is derived from residual tissue autoflourescence and common staining noise in the background results from nonspecific antibody deposition secondary to necrosis, tissue folds and fixation variations. MIBI has its own unique challenges for denoising due to low intensity values and sparse and pixelated signal for low abundant antigens (1).

The MIBI dataset was acquired from bladder, breast and lung cancer patients and the cell nuclei in the double-stranded DNA (dsDNA) channel were semi-automatically segmented and manually-corrected by a trained technician. These datasets consist of single channel patches of size 512×512. The breast cancer dataset contained 160 patches from 25 acquisitions, the bladder cancer dataset 110 patches from 30 acquisitions, and the lung cancer dataset 120 patches from 30 acquisitions.

The remaining datasets were manually annotated by a trained technician. The image patches have a reduced 400×400 size, and consisted of a single dapi (nuclei) channel Vectra dataset containing 36 annotated image patches, 8 single dsDNA (nuclei) channel MIBI image patches, 10 2-channel (dapi + PanCK) Vectra patches, and 4 2-channel (dsDNA + PanCK) MIBI image patches.

## Results

We compared the performance of our method in both semantic and instance segmentation against three models. A baseline model (Base) that implements the *blind-spot network* training scheme and only trains on the segmentation loss on the available scribbles. We also compared against DenoiSeg (DSeg) (28), and Scribble2Label (S2L) (23). Since DSeg builds on the Noise2Void framework, we trained their model with the same U-Net architecture and *blind-spot network* specifications as ours. S2L was trained under different configurations of network architecture, and hyper-parameters (details provided in supplementary material), the best result is reported. All methods were trained until convergence with a stopping criteria on validation loss, using a batch size of 64 unless otherwise specified. We report semantic segmentation performance with *F*1 score and AUC metrics, and instance segmentation with mean intersection over union (mIoU) and mean Dice-coefficient (mDice). In all cases, we report performance for varying number of instance scribbles (*n_s_)*, we focused on the low-annotation regime.

### Preprocessing

We split each dataset into a set of training and test images; all images are percentile normalized in the range [0.1,99.8]%. For a given training image set and instance scribble budget *n_s_*, we generate random scribbles from the available ground truth masks. We then sample the training images and corresponding scribbles into patches of size 128 x 128 using the tools provided in (30). Validation patches are sampled from the same training set, but we ensure they do not overlap more than a fixed percentage with the training patches. The training patches were augmented following the 8-fold data augmentation used in (28), which consisted on 90 degree rotations and flipping of all images. All methods were trained over the same training and validation dataset patches, without any additional augmentation. Since DSeg requires boundary labels, their method was given additional information on whether a background scribble was part of an instance boundary.

As a point of reference for wall-clock algorithm time, on the MIBI-Manual dataset, a batch of 64 images can be processed in 0.21 seconds on the S2L implementation, 1.18 seconds on DSeg, and 1.11 on ImPartial. We note that the time gains made on S2L are partly due to their implementation using 8 parallel workers, while both DSeg and ImPartial are currently built on top of CARE on tensor flow.

### MIBI Modality

We evaluate the performance of the proposed method on single dsDNA channel images for breast, bladder and lung cancer patients datasets with semi-automatic ground truth labels. For each dataset, we randomly generate a budget of instance scribbles (*n_s_)* for image patches corresponding to 10 of the available acquisitions, the remaining patches are consider as test examples. In Table 1, we report the performance on semantic and instance segmentation on the train and test examples. We indicate the number of instance scribbles provided (*n_s_)* and the number of total instances (*n_i_*). Figure 3 shows example segmentation results produced by the tested methods.

**Fig. 3.**
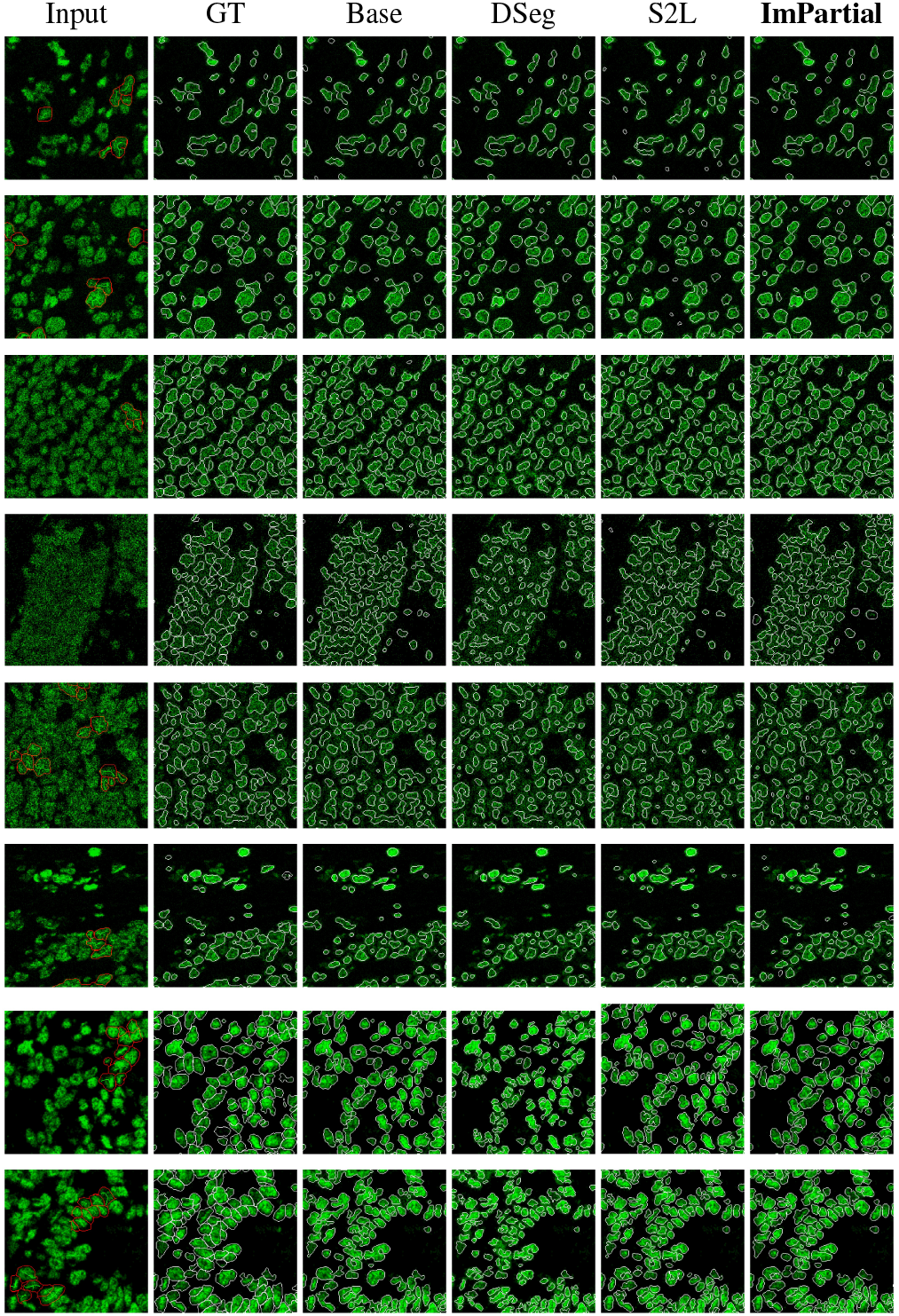
Here we show MIBI Bladder (rows 1&2), Breast (rows 3&4), Lung (rows 5&6) and MIBI-Manual (rows 7&8) segmentation examples for each method. First column shows the input image and the available scribbles. Note the variability of nuclei structures across the different samples

**Table 1.** Comparison of semantic and instance segmentation performance of the evaluated methods on MIBI 1-Channel (dsDNA) datasets. We indicate the number of annotated instance scribbles (ns) versus the total number of available instances (ni). We also report the performance on test data not previously seen by the model.

| Semantic and Instance segmentation MIBI Breast |  |  |  |  |  |  |  |  |  |  |  |  |  |  |  |  |
| --- | --- | --- | --- | --- | --- | --- | --- | --- | --- | --- | --- | --- | --- | --- | --- | --- |
|  | Base | DSeg | S2L | Ours | Base | DSeg | S2L | Ours | Base | DSeg | S2L | Ours | Base | DSeg | S2L | Ours |
| ns/ni | F1 score |  |  |  | AUC score |  |  |  | mDICE score |  |  |  | mIOU score |  |  |  |
| 250/25.5k | 0.766 | 0.796 | 0.755 | **0.799** | 0.952 | 0.949 | 0.962 | **0.973** | 0.74 | 0.685 | 0.715 | **0.748** | 0.627 | 0.577 | 0.599 | **0.637** |
| 0/15.6k (test) | 0.596 | 0.686 | 0.555 | **0.717** | 0.967 | 0.96 | 0.972 | **0.986** | 0.734 | 0.691 | **0.74** | **0.74** | 0.61 | 0.571 | 0.609 | **0.618** |
| 500/25.5k | 0.796 | 0.801 | 0.76 | **0.822** | 0.967 | 0.971 | 0.953 | **0.974** | 0.754 | 0.704 | 0.745 | **0.762** | 0.648 | 0.604 | 0.634 | **0.657** |
| 0/15.6k (test) | 0.68 | 0.701 | 0.672 | **0.795** | 0.982 | **0.988** | 0.964 | 0.987 | 0.746 | 0.735 | 0.738 | **0.753** | 0.64 | 0.643 | 0.631 | **0.646** |
| 1k/25.5k | 0.795 | 0.813 | 0.789 | **0.83** | 0.965 | 0.967 | 0.973 | **0.979** | 0.758 | 0.732 | 0.753 | **0.775** | 0.649 | 0.623 | 0.644 | **0.657** |
| 0/15.6k (test) | 0.731 | 0.754 | 0.665 | **0.786** | 0.975 | **0.987** | 0.984 | 0.982 | 0.737 | 0.782 | 0.752 | **0.796** | 0.622 | 0.673 | 0.626 | **0.684** |
| Semantic and Instance segmentation MIBI Bladder |  |  |  |  |  |  |  |  |  |  |  |  |  |  |  |  |
|  | Base | DSeg | S2L | Ours | Base | DSeg | S2L | Ours | Base | DSeg | S2L | Ours | Base | DSeg | S2L | Ours |
| ns/ni | F1 score |  |  |  | AUC score |  |  |  | mDICE score |  |  |  | mIOU score |  |  |  |
| 250/7.6k | 0.779 | **0.82** | 0.712 | 0.774 | 0.978 | **0.981** | 0.946 | 0.975 | 0.785 | 0.775 | 0.759 | **0.793** | 0.682 | 0.673 | 0.649 | **0.688** |
| 0/18.7k (test) | 0.789 | **0.801** | 0.715 | 0.784 | 0.974 | **0.977** | 0.939 | 0.971 | 0.751 | 0.745 | 0.744 | **0.765** | 0.644 | 0.64 | 0.632 | **0.656** |
| 500/7.6k | 0.732 | **0.824** | 0.729 | 0.787 | 0.954 | 0.968 | 0.959 | **0.973** | 0.811 | 0.767 | 0.806 | **0.815** | 0.718 | 0.673 | 0.707 | **0.736** |
| 0/18.7k (test) | 0.749 | **0.825** | 0.735 | 0.781 | 0.95 | 0.965 | 0.953 | **0.966** | 0.786 | 0.758 | 0.787 | **0.799** | 0.688 | 0.659 | 0.682 | **0.705** |
| 1k/7.6k | 0.833 | 0.83 | 0.769 | **0.839** | 0.987 | 0.984 | 0.975 | **0.988** | 0.816 | 0.804 | 0.8 | **0.82** | 0.721 | 0.708 | 0.701 | **0.74** |
| 0/18.7k (test) | 0.821 | 0.817 | 0.782 | **0.826** | 0.986 | 0.981 | 0.971 | **0.988** | 0.795 | 0.783 | 0.781 | **0.802** | 0.704 | 0.682 | 0.676 | **0.712** |
| Semantic and Instance segmentation MIBI Lung |  |  |  |  |  |  |  |  |  |  |  |  |  |  |  |  |
|  | Base | DSeg | S2L | Ours | Base | DSeg | S2L | Ours | Base | DSeg | S2L | Ours | Base | DSeg | S2L | Ours |
| ns/ni | F1 score |  |  |  | AUC score |  |  |  | mDICE score |  |  |  | mIOU score |  |  |  |
| 250/8.4k | 0.738 | 0.766 | 0.75 | **0.781** | 0.964 | 0.963 | 0.969 | **0.978** | 0.723 | 0.711 | 0.736 | **0.753** | 0.612 | 0.613 | 0.626 | **0.654** |
| 0/13.1k (test) | 0.69 | 0.715 | 0.693 | **0.734** | 0.936 | 0.925 | 0.94 | **0.953** | 0.666 | 0.656 | **0.697** | **0.697** | 0.561 | 0.564 | 0.592 | **0.605** |
| 500/8.4k | 0.763 | **0.78** | 0.751 | 0.774 | **0.965** | 0.948 | 0.956 | 0.962 | 0.704 | 0.704 | **0.731** | 0.722 | 0.606 | 0.608 | **0.627** | 0.622 |
| 0/13.1k (test) | 0.714 | 0.725 | 0.677 | **0.726** | **0.941** | 0.916 | 0.926 | 0.936 | 0.65 | 0.641 | 0.66 | **0.669** | 0.554 | 0.551 | 0.571 | **0.574** |
| 1k/8.4k | 0.769 | **0.782** | 0.739 | 0.769 | **0.977** | 0.967 | 0.958 | 0.974 | 0.742 | 0.748 | 0.736 | **0.763** | 0.64 | 0.649 | 0.63 | **0.657** |
| 0/13.1k (test) | 0.724 | **0.732** | 0.686 | 0.722 | **0.949** | 0.927 | 0.925 | 0.945 | 0.701 | 0.693 | 0.702 | **0.72** | 0.611 | 0.598 | 0.6 | **0.62** |
| Semantic and Instance segmentation MIBI-Manual |  |  |  |  |  |  |  |  |  |  |  |  |  |  |  |  |
|  | Base | DSeg | S2L | Ours | Base | DSeg | S2L | Ours | Base | DSeg | S2L | Ours | Base | DSeg | S2L | Ours |
| ns/ni | F1 score |  |  |  | AUC score |  |  |  | mDICE score |  |  |  | mIOU score |  |  |  |
| 50/893 | **0.876** | 0.702 | 0.841 | 0.864 | **0.974** | 0.832 | 0.957 | 0.968 | 0.631 | 0.628 | 0.698 | **0.725** | 0.514 | 0.503 | 0.58 | **0.609** |
| 0/812 (test) | **0.866** | 0.696 | 0.835 | 0.853 | **0.98** | 0.816 | 0.966 | 0.974 | 0.642 | 0.652 | 0.709 | **0.732** | 0.53 | 0.512 | 0.591 | **0.618** |
| 100/893 | 0.818 | 0.723 | 0.817 | **0.844** | **0.938** | 0.821 | 0.935 | 0.933 | **0.698** | 0.656 | 0.695 | 0.695 | **0.583** | 0.538 | 0.574 | **0.583** |
| 0/812 (test) | 0.813 | 0.721 | 0.809 | **0.838** | **0.947** | 0.81 | 0.94 | 0.941 | 0.71 | 0.668 | 0.705 | **0.712** | 0.601 | 0.554 | 0.584 | **0.604** |
| 200/893 | 0.835 | 0.743 | **0.843** | 0.829 | **0.961** | 0.833 | 0.953 | 0.957 | 0.715 | 0.719 | **0.721** | 0.719 | 0.6 | 0.602 | **0.611** | 0.603 |
| 0/812 (test) | 0.826 | **0.836** | **0.836** | 0.822 | **0.966** | 0.958 | 0.958 | 0.962 | 0.729 | 0.729 | 0.728 | **0.731** | 0.617 | **0.62** | 0.615 | 0.617 |

On the MIBI single dsDNA channel dataset with manual annotations (MIBI-Manual), we considered 4 image patches to provide instance scribbles and the remaining 4 for test. For the MIBI 2-channel manual annotated dataset (MIBI 2- Channels), we considered 2 image patches for training and 2 for testing. Quantitative results are provided in tables 1 and 2, visuals of the MIBI 2-Channels segmentation are presented in Figure 4.

**Fig. 4.**
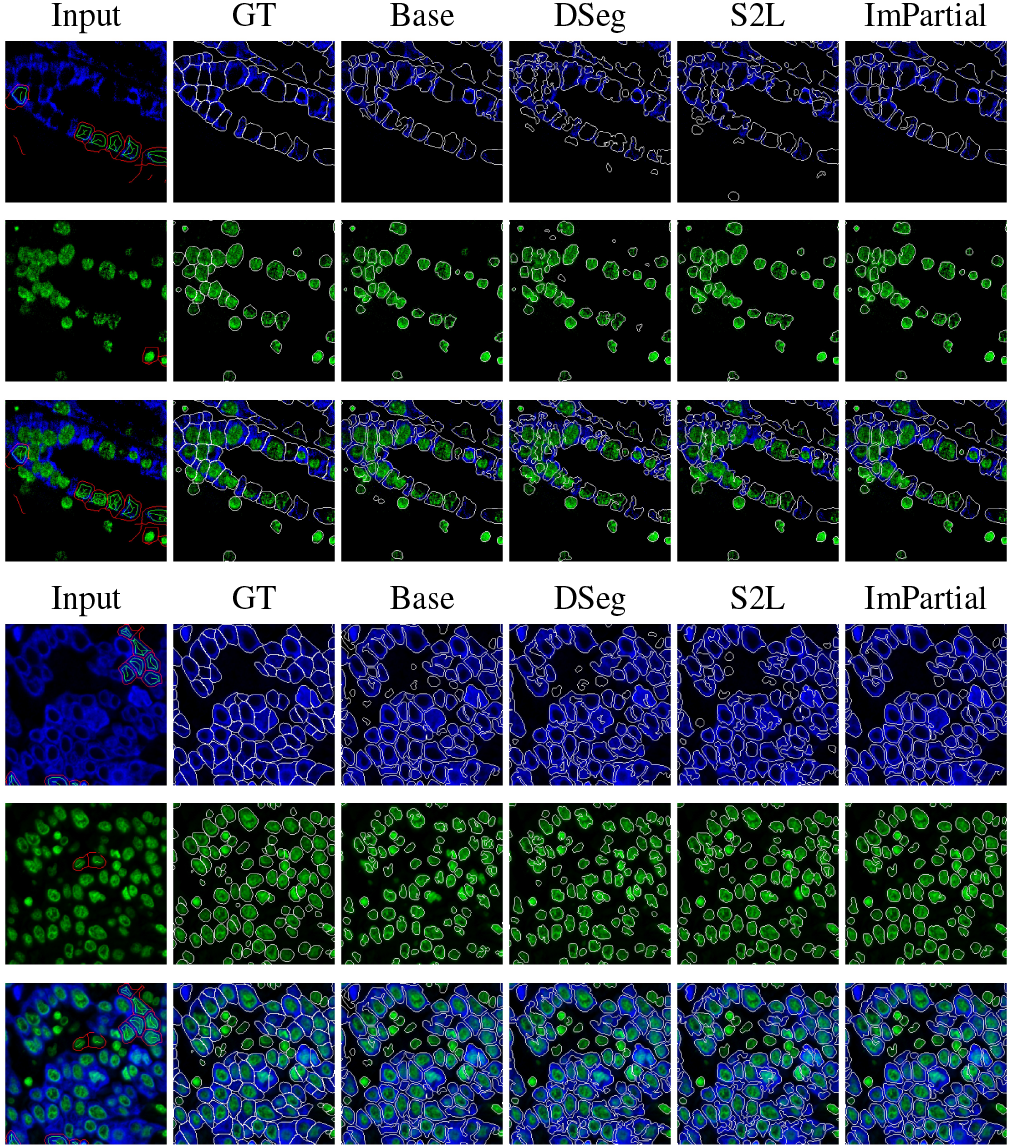
Comparison of the instance segmentation mask recovered by every considered method on a two-channel MIBI and Vectra image. First three rows correspond to the cytoplasm channel (PanCK), the nuclei channel (dsDNA), and composite image (cytoplasm and nuclei channels) for the MIBI modality. Last three rows correspond to the cytoplasm channel (PanCK), the nuclei channel (dapi), and composite image (cytoplasm and nuclei channels) for the Vectra modality. The scribbles for the desired semantic segmentation are also shown, models were trained with 100 cytoplasm and 100 nuclei instance scribbles. Second column contains ground truth segmentation for each of the semantic masks (all nuclei and cytoplasm, nuclei surrounded by cytoplasm, and all nuclei respectively). Following columns show the contour of the instances recovered by the baseline model (network without mixture loss), DSeg, S2L and ImPartial.

**Table 2.** Comparison of semantic and instance segmentation performance of the evaluated methods on the MIBI 2 Channels (PanCK and dsDNA) dataset. We indicate the number of annotated instance scribbles (ns) versus the total number of available instances (ni). We also report the performance on test data not previously seen by the model.

| Semantic and Instance segmentation MIBI PanCK |  |  |  |  |  |  |  |  |  |  |  |  |  |  |  |  |
| --- | --- | --- | --- | --- | --- | --- | --- | --- | --- | --- | --- | --- | --- | --- | --- | --- |
|  | Base | DSeg | S2L | Ours | Base | DSeg | S2L | Ours | Base | DSeg | S2L | Ours | Base | DSeg | S2L | Ours |
| ns/ni | F1 score |  |  |  | AUC score |  |  |  | mDice score |  |  |  | mIOU score |  |  |  |
| 25/234 | **0.838** | 0.81 | 0.807 | **0.838** | 0.882 | 0.879 | 0.915 | **0.935** | 0.604 | 0.499 | 0.554 | **0.61** | 0.475 | 0.38 | 0.422 | **0.476** |
| 0/111 (test) | 0.611 | 0.548 | 0.49 | **0.62** | 0.84 | 0.861 | 0.853 | **0.913** | 0.51 | 0.428 | 0.45 | **0.574** | 0.384 | 0.324 | 0.318 | **0.443** |
| 50/234 | **0.866** | 0.848 | 0.812 | 0.863 | **0.928** | 0.912 | 0.915 | 0.925 | 0.669 | 0.559 | 0.61 | **0.67** | **0.546** | 0.436 | 0.481 | 0.543 |
| 0/111 (test) | **0.746** | 0.617 | 0.472 | 0.744 | 0.905 | 0.89 | 0.831 | **0.907** | 0.446 | 0.425 | **0.535** | 0.519 | 0.324 | 0.315 | **0.398** | 0.396 |
| 100/234 | **0.879** | 0.864 | 0.829 | 0.876 | 0.928 | 0.943 | 0.928 | **0.949** | 0.714 | 0.648 | 0.642 | **0.719** | 0.592 | 0.517 | 0.513 | **0.599** |
| 0/111 (test) | 0.669 | 0.647 | 0.513 | **0.686** | 0.88 | 0.915 | 0.845 | **0.917** | 0.528 | 0.46 | 0.525 | **0.599** | 0.404 | 0.351 | 0.393 | **0.467** |
| Semantic and Instance segmentation MIBI dsDNA |  |  |  |  |  |  |  |  |  |  |  |  |  |  |  |  |
|  | Base | DSeg | S2L | Ours | Base | DSeg | S2L | Ours | Base | DSeg | S2L | Ours | Base | DSeg | S2L | Ours |
| ns/ni | F1 score |  |  |  | AUC score |  |  |  | mDice score |  |  |  | mIOU score |  |  |  |
| 25/215 | 0.767 | **0.842** | 0.766 | 0.817 | **0.937** | 0.96 | 0.932 | 0.934 | 0.71 | 0.656 | 0.636 | **0.749** | 0.59 | 0.551 | 0.513 | **0.637** |
| 0/270 (test) | 0.784 | **0.839** | 0.781 | 0.824 | 0.958 | **0.96** | 0.936 | 0.933 | 0.683 | 0.625 | 0.592 | **0.723** | 0.571 | 0.516 | 0.487 | **0.615** |
| 50/215 | 0.789 | **0.869** | 0.792 | 0.8 | 0.958 | **0.974** | 0.941 | 0.935 | 0.756 | 0.699 | 0.733 | **0.765** | 0.638 | 0.587 | 0.618 | **0.651** |
| 0/270 (test) | 0.748 | **0.836** | 0.745 | 0.75 | 0.942 | **0.964** | 0.921 | 0.92 | 0.696 | 0.664 | 0.687 | **0.707** | 0.579 | 0.557 | 0.567 | **0.584** |
| 100/215 | 0.841 | **0.858** | 0.716 | 0.837 | 0.946 | 0.958 | 0.947 | **0.966** | **0.777** | 0.731 | 0.693 | 0.77 | 0.657 | 0.619 | 0.562 | **0.669** |
| 0/270 (test) | 0.819 | **0.842** | 0.663 | 0.811 | 0.957 | 0.944 | 0.936 | **0.958** | 0.752 | 0.716 | 0.646 | **0.759** | 0.645 | 0.596 | 0.521 | **0.648** |
| Semantic and Instance segmentation MIBI joint PanCK + dsDNA |  |  |  |  |  |  |  |  |  |  |  |  |  |  |  |  |
|  | Base | DSeg | S2L | Ours | Base | DSeg | S2L | Ours | Base | DSeg | S2L | Ours | Base | DSeg | S2L | Ours |
| ns/ni | F1 score |  |  |  | AUC score |  |  |  | mDice score |  |  |  | mIOU score |  |  |  |
| 25/239 | 0.848 | 0.845 | **0.855** | 0.846 | 0.897 | 0.941 | 0.939 | **0.943** | 0.659 | 0.584 | 0.586 | **0.663** | 0.539 | 0.461 | 0.463 | **0.54** |
| 0/262 (test) | 0.825 | **0.842** | 0.727 | 0.813 | 0.917 | **0.974** | 0.958 | 0.962 | 0.651 | 0.586 | 0.586 | **0.668** | 0.536 | 0.477 | 0.44 | **0.548** |
| 50/239 | 0.841 | **0.866** | 0.848 | 0.853 | 0.926 | 0.931 | **0.939** | 0.918 | 0.677 | 0.574 | 0.64 | **0.697** | 0.562 | 0.452 | 0.515 | **0.576** |
| 0/262 (test) | 0.77 | **0.831** | 0.675 | 0.774 | 0.914 | **0.955** | 0.91 | 0.907 | 0.525 | 0.494 | 0.526 | **0.571** | 0.422 | 0.372 | 0.439 | **0.446** |
| 100/239 | **0.874** | 0.85 | 0.859 | 0.87 | 0.928 | 0.928 | 0.945 | **0.949** | 0.719 | 0.648 | 0.647 | **0.724** | 0.597 | 0.533 | 0.522 | **0.609** |
| 0/262 (test) | 0.821 | **0.842** | 0.675 | 0.819 | 0.921 | 0.944 | 0.936 | **0.954** | 0.635 | **0.676** | 0.647 | 0.658 | 0.513 | **0.586** | 0.54 | 0.536 |

We note that results on both semantic and instance segmentation produced by ImPartial outperform previous methods in many cases. Qualitatively, we observe the segmentation recovered by our method has less blocky artifacts and better matches the overall shape and structure of the underlying cells and cytoplasm instances.

### Vectra Modality

We evaluate the performance of the proposed method on two Vectra datasets. The first one consisted of a single channel showing cell nuclei (dapi), results are presented in Table 3. The second one shows a two-channel cytoplasm and nuclei (dapi + PanCK) acquisition; here the goal was to simultaneously segment instances of cytoplasm (blue channel in 4), instances of nuclei (green channel in 4) and a combination of both (simulating chromogenic bright-field 2-channel acquisition). Results are presented in Table 4. For the single dapi channel dataset, 10 patches were selected for training; their ground truth annotations were used to randomly generate a budget of instance scribbles (*n_s_).* The remaining 26 patches were used as test examples. For the 2-channel dapi+PanCK dataset, we used 4 patches as training examples; instance scribbles were randomly generated for dapi (nuclei), PanCK (cytoplasm) and a combination of both. The remaining 4 patches were set as test examples. On this last dataset, all methods used a U-net with depth 2 and an initial filter bank of 128 feature maps, batch size was set to 32 due to GPU capacity. DSeg and S2L implementations do not natively handle multiple classification objectives, so they were trained on each objective individually, using the 2-channel input.

**Table 3.** Comparison of semantic and instance segmentation performance of the evaluated methods on the Vectra 1-channel nuclei segmentation dataset. We indicate the number of annotated instance scribbles (ns) versus the total number of available instances (ni). We also report the performance on test data not previously seen by the model.

|  | Base | DSeg | S2L | Ours | Base | DSeg | S2L | Ours | Base | DSeg | S2L | Ours | Base | DSeg | S2L | Ours |
| --- | --- | --- | --- | --- | --- | --- | --- | --- | --- | --- | --- | --- | --- | --- | --- | --- |
| ns/ni | F1 score |  |  |  | AUC score |  |  |  | mDice score |  |  |  | mIOU score |  |  |  |
| 50/4.7k | **0.75** | 0.74 | 0.692 | 0.748 | 0.877 | 0.878 | 0.834 | **0.893** | 0.621 | 0.52 | 0.604 | **0.63** | 0.491 | 0.389 | 0.471 | **0.494** |
| 0/11.4k (test) | 0.728 | 0.723 | 0.681 | **0.73** | 0.854 | 0.86 | 0.821 | **0.88** | 0.619 | 0.521 | 0.595 | **0.628** | 0.49 | 0.387 | 0.461 | **0.493** |
| 100/4.7k | 0.748 | 0.755 | 0.743 | **0.774** | 0.877 | 0.859 | 0.882 | **0.901** | **0.666** | 0.571 | 0.628 | 0.665 | **0.536** | 0.439 | 0.499 | 0.534 |
| 0/11.4k (test) | 0.718 | 0.737 | 0.725 | **0.745** | 0.855 | 0.833 | 0.864 | **0.885** | 0.637 | 0.552 | 0.6 | **0.639** | 0.502 | 0.419 | 0.468 | **0.503** |
| 250/4.7k | 0.77 | 0.771 | **0.778** | **0.778** | **0.915** | 0.864 | 0.897 | **0.915** | 0.659 | 0.611 | 0.639 | **0.662** | 0.527 | 0.479 | 0.51 | **0.534** |
| 0/11.4k (test) | 0.745 | 0.744 | 0.752 | **0.755** | **0.906** | 0.835 | 0.879 | 0.905 | 0.633 | 0.584 | 0.619 | **0.64** | 0.499 | 0.449 | 0.486 | **0.508** |

**Table 4.** Comparison of semantic and instance segmentation performance of the evaluated methods on the Vectra 2 Channels (PanCK and dapi) dataset. We indicate the number of annotated instance scribbles (ns) versus the total number of available instances (ni). We also report the performance on test data not previously seen by the model.

| Semantic and Instance segmentation Vectra PanCK |  |  |  |  |  |  |  |  |  |  |  |  |  |  |  |  |
| --- | --- | --- | --- | --- | --- | --- | --- | --- | --- | --- | --- | --- | --- | --- | --- | --- |
|  | Base | DSeg | S2L | Ours | Base | DSeg | S2L | Ours | Base | DSeg | S2L | Ours | Base | DSeg | S2L | Ours |
| ns/ni | F1 score |  |  |  | AUC score |  |  |  | mDICE score |  |  |  | mIOU score |  |  |  |
| 50/961 | 0.83 | 0.832 | **0.848** | 0.828 | 0.898 | 0.877 | 0.889 | **0.891** | 0.501 | 0.477 | 0.495 | **0.534** | 0.393 | 0.361 | 0.381 | **0.423** |
| 0/699 (test) | 0.826 | 0.77 | 0.802 | **0.827** | **0.943** | 0.919 | 0.933 | 0.934 | 0.632 | 0.564 | 0.581 | **0.652** | 0.519 | 0.444 | 0.462 | **0.54** |
| 100/961 | 0.813 | 0.775 | **0.842** | 0.824 | 0.898 | 0.821 | 0.892 | **0.912** | 0.628 | 0.536 | 0.534 | **0.63** | 0.505 | 0.414 | 0.416 | **0.512** |
| 0/699 (test) | 0.79 | 0.697 | **0.818** | 0.801 | 0.914 | 0.837 | **0.946** | 0.932 | 0.678 | 0.589 | 0.617 | **0.685** | 0.569 | 0.458 | 0.498 | **0.576** |
| 250/961 | 0.85 | 0.84 | 0.858 | **0.859** | 0.93 | 0.887 | 0.926 | **0.932** | 0.658 | 0.589 | 0.606 | **0.67** | **0.547** | 0.467 | 0.485 | 0.537 |
| 0/699 (test) | 0.827 | 0.823 | **0.839** | 0.831 | 0.938 | 0.911 | 0.961 | **0.945** | 0.695 | 0.627 | 0.687 | **0.703** | 0.586 | 0.512 | 0.573 | **0.593** |
| Semantic and Instance segmentation Vectra dapi |  |  |  |  |  |  |  |  |  |  |  |  |  |  |  |  |
|  | Base | DSeg | S2L | Ours | Base | DSeg | S2L | Ours | Base | DSeg | S2L | Ours | Base | DSeg | S2L | Ours |
| ns/ni | F1 score |  |  |  | AUC score |  |  |  | mDICE score |  |  |  | mIOU score |  |  |  |
| 50/918 | 0.807 | 0.838 | 0.751 | **0.841** | 0.933 | **0.946** | 0.901 | 0.932 | 0.675 | 0.607 | 0.617 | **0.682** | 0.557 | 0.49 | 0.496 | **0.569** |
| 0/1.2k (test) | 0.802 | 0.829 | 0.719 | **0.836** | 0.946 | 0.951 | 0.887 | **0.957** | **0.729** | 0.666 | 0.66 | 0.728 | 0.617 | 0.553 | 0.54 | **0.622** |
| 100/918 | 0.773 | 0.84 | 0.767 | **0.825** | 0.932 | **0.965** | 0.917 | 0.946 | 0.664 | 0.579 | 0.613 | **0.68** | 0.543 | 0.465 | 0.496 | **0.566** |
| 0/1.2k (test) | 0.763 | **0.841** | 0.751 | 0.825 | 0.94 | **0.976** | 0.913 | 0.964 | 0.695 | 0.653 | 0.678 | **0.73** | 0.575 | 0.546 | 0.563 | **0.622** |
| 250/918 | 0.799 | 0.807 | 0.825 | **0.834** | 0.951 | 0.919 | 0.954 | **0.963** | 0.676 | 0.64 | 0.626 | **0.684** | 0.558 | 0.521 | 0.507 | **0.568** |
| 0/1.2k (test) | 0.784 | 0.803 | 0.766 | **0.826** | 0.962 | 0.938 | 0.938 | **0.973** | 0.714 | 0.686 | 0.671 | **0.743** | 0.598 | 0.582 | 0.55 | **0.605** |
| Semantic and Instance segmentation Vectra joint PanCK + dapi |  |  |  |  |  |  |  |  |  |  |  |  |  |  |  |  |
|  | Base | DSeg | S2L | Ours | Base | DSeg | S2L | Ours | Base | DSeg | S2L | Ours | Base | DSeg | S2L | ImPartial |
| ns/ni | F1 score |  |  |  | AUC score |  |  |  | mDICE score |  |  |  | mIOU score |  |  |  |
| 50/918 | 0.835 | 0.83 | **0.85** | 0.834 | **0.897** | 0.874 | 0.886 | 0.875 | 0.496 | 0.432 | 0.455 | **0.552** | 0.384 | 0.333 | 0.351 | **0.434** |
| 0/1.23k (test) | 0.819 | 0.809 | 0.751 | **0.823** | **0.938** | 0.922 | 0.905 | 0.91 | 0.566 | 0.458 | 0.61 | **0.645** | 0.456 | 0.36 | 0.485 | **0.529** |
| 100/918 | 0.809 | 0.811 | **0.841** | 0.83 | 0.9 | 0.871 | 0.904 | **0.906** | **0.635** | 0.537 | 0.559 | 0.631 | **0.514** | 0.418 | 0.44 | 0.511 |
| 0/1.23k (test) | 0.802 | 0.808 | 0.762 | **0.809** | 0.921 | 0.916 | 0.896 | **0.929** | 0.645 | 0.546 | 0.645 | **0.647** | 0.527 | 0.429 | 0.528 | **0.531** |
| 250/918 | 0.844 | 0.838 | **0.859** | 0.844 | 0.926 | 0.894 | 0.917 | **0.929** | 0.629 | 0.576 | 0.561 | **0.654** | 0.511 | 0.459 | 0.446 | **0.534** |
| 0/1.23k (test) | **0.816** | 0.81 | 0.768 | **0.816** | 0.939 | 0.906 | 0.916 | **0.946** | 0.66 | 0.561 | 0.616 | **0.684** | 0.547 | 0.448 | 0.51 | **0.568** |

We achieved superior performance on the instance and semantic segmentation metrics. A similar phenomenon to the MIBI results was observed, where the recovered instances had shapes more closely resembling ground truth labels, with less spurious detections and segmentation noise. Our method was also able to accomplish 3 different segmentation tasks on the 2-channel Vectra dataset without any further adaptations, which shows that it is able to adequately process multichannel and multi-objective data. It is worth noting that on both the Vectra and MIBI datasets, the blind-spot network shows remarkably good performance, which may indicate that, in these settings, there is limited need for a large number of annotations to get a decent baseline performance. This latter observation is also supported by the relatively minor gains in performance we observe when we increase the number of scribbles available to the training agent.

### Conclusion and Future Work

Here we proposed a weakly-supervised method to segment nuclei from noisy images when only few scribble annotations of background and foreground are provided. We compared the performance of our method and recently proposed techniques on a variety of biological datasets, showing that in most cases our approach achieves the best performance on instance segmentation. The analysis presented in this paper shows that all methods achieve a decent baseline performance without a large number of annotations. Moreover, increasing the number of annotations does not necessarily translate to a large gain in performance.

Future work will analyze the use of continual learning techniques to develop a more interactive system where the user can improve on the segmentation at any given moment by iteratively providing more scribbles. This will require leveraging techniques from backwards compatibility to ensure correct segmentation are not lost in the updating process. From our observations we think it is worth analyzing how can the quality of the manual annotations be improved in order to obtain quality samples and reduce the amount of unnecessary annotations’

In future work, we plan to develop a Web User Interface where people can upload their dataset, provide scribble annotations for a few examples and download the corresponding segmentations’ A related line of work towards the goal of annotation-as-a-service includes visualizing out-of-distribution areas of the image so that the user can be especially attentive for potential defects in the segmentation in those areas’

## ACKNOWLEDGEMENTS

This study was supported by MSK Cancer Center Support Grant/Core Grant (P30 CA008748).

## Footnotes

1 https://www.akoyabio.com/phenoptics/mantra-vectra-instruments/vectra-polaris/

2 https://www.ionpath.com/mibi-technology/

